# Molecular mechanisms of gating in the calcium-activated chloride channel bestrophin

**DOI:** 10.1101/432682

**Authors:** Alexandria N. Miller, George Vaisey, Stephen B. Long

## Abstract

Bestrophin (BEST1–4 in humans) channels are ligand gated chloride (Cl^−^) channels that are activated by calcium (Ca^2+^). Mutations in BEST1 cause retinal degenerative diseases. Partly because these channels have no sequence or structural similarity to other ion channels, the molecular mechanisms underlying gating are unknown. Here, we present a series of cryo-electron microscopy (cryo-EM) structures of chicken BEST1, determined at 3.1 Å resolution or better, that represent the principal gating states of the channel. Unlike other channels, opening of the pore is due to the repositioning of tethered pore-lining helices within a surrounding protein shell that dramatically widens a “neck” of the pore through a concertina of amino acid rearrangements within the protein core. The neck serves as both the activation and the inactivation gate. The binding of Ca^2+^ to a cytosolic domain instigates pore opening and the structures reveal that, unlike voltage-gated Na^+^ and K^+^ channels, similar molecular rearrangements are responsible for inactivation and deactivation. A single aperture within the 95 Å-long opened pore separates the cytosol from the extracellular milieu and controls anion permeability. The studies define the basis for Ca^2+^-activated Cl^−^ channel function and reveal a new molecular paradigm for gating in ligand-gated ion channels.

---

The family of bestrophin proteins (BEST1–4) was identified by linkage analysis to hereditary macular degenerations caused by mutations in BEST1 (*1, 2*); to date more than 200 mutations in BEST1 are linked with eye disease (*3, 4*). BEST1–4 proteins are expressed in the plasma membrane and form Ca^2+^-activated Cl^−^ channels by assembling as pentamers (*5–9*). Data suggest that BEST1 mediates a Ca^2+^-activated Cl^−^ current that is integral to human retinal pigment epithelial function (*10*). The broad tissue distribution of the family suggests additional physiological functions that are not fully realized (*5*), and these may include processes as diverse as cell volume regulation (*11*), pH homeostasis (*12*), and neurotransmitter release (*13*). An X-ray structure of chicken BEST1, which shares 74 % sequence identity with human BEST1 and possesses analogous Ca^2+^-activation and anion-selectivity properties, revealed an architecture that is distinct from other ion channel families (*6*). A prokaryotic ion channel (KpBest) has discernable sequence (14 % identity) and structural homology with BEST1, but the Ca^2+^-activated Cl^−^ channel function of bestrophin proteins appears specific to metazoan organisms; KpBest is cation-selective and not activated by Ca^2+^ (*14*). The pore of BEST1 contains two hydrophobic constrictions, the “neck” and the “aperture”, and ostensibly either of these could function as a gate. Ionic currents through BEST1 have recently been found to decrease over time due to the inactivation of the channel, which is caused by the binding of a C-terminal peptide to a receptor on the channel’s cytosolic surface (*15*). Because the inactivation peptide is bound to its receptor in the structure and because the antibody that was used as a crystallization chaperone also promotes inactivation, we now realize that the previously determined structure of BEST1 likely represents an inactivated state (6, 15). How the channel opens is not known.

To begin to address the conformational changes associated with channel opening, wedetermined a 3.1 Å resolution cryo-EM structure of BEST1 without an antibody (the same construct used for X-ray studies, comprising residues 1–405, and termed BEST1_405_). From single particle analysis we obtained a single conformation, which is indistinguishable from the X-ray structure, including a bound inactivation peptide, and therefore this presumably also represents a Ca^2+^-bound inactivated state (fig. S1, fig. S3; RMSD for Cα atoms =0.5 Å). This result suggests that antibody binding does not distort the channel from a native conformation and that differences we might observe in channel structures are not the result of differences in methodologies between cryo-EM and X-ray crystallography.

In an aim to obtain structural information on an open conformation of the channel, we removed the C-terminal inactivation peptide (by using a construct spanning amino acids 1–345, termed BEST1_345_) and used this for cryo-EM studies. Importantly, whilst BEST1_345_ does not inactivate, it possesses normal Cl^−^ selectivity and Ca^2+^-dependent activation (*15*). Single particle cryo-EM analysis of Ca^2+^-bound BEST1_345_ (fig. S2, fig. S3, fig. S5) revealed two distinct conformations of the channel (Fig. 1). The first, determined at 3.0 Å resolution, represents 86% of the particles and is essentially indistinguishable from the structure of BEST1_405_ (Fig. 1a, fig. S1c, RMSD for Cα atoms =0.5 Å), except for the absence of the inactivation peptide. Because BEST1_345_ does not inactivate, the structure presumably represents a Ca^2+^-bound closed conformation. In this and in the inactivated structure, the neck adopts an indistinguishable closed conformation (Fig. 2c). The second structure, which represents 14% of the particles and is determined to 2.9 Å resolution, contains a dramatically widened pore within the neck (Fig. 1b). Based on discussions presented herein we conclude that this represents the open conformation of the channel. The relative abundance of the closed conformation suggests that it is energetically favorable.

**Fig. 1.**
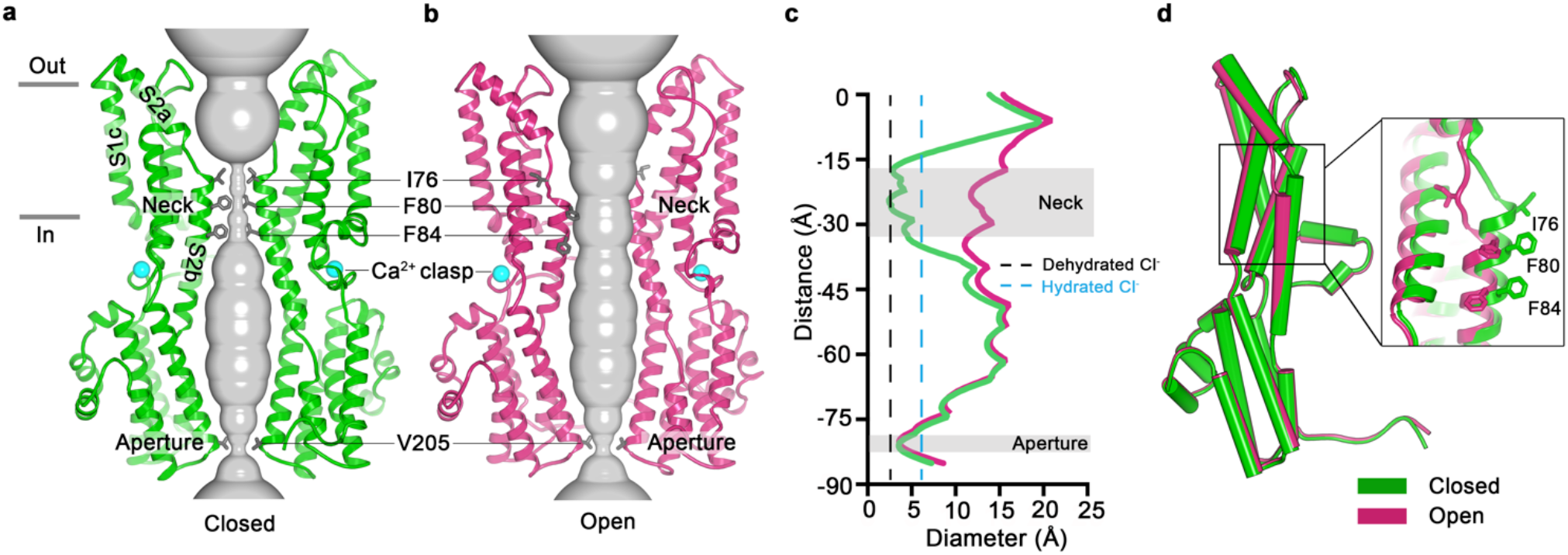
Open and closed conformations. **a-b**, Cutaway views of the Ca^2+^-bound closed (green; nonconductive) and Ca^2+^-bound open (pink) conformations of BEST1_345_. The minimal radial distance from the center of the pore to the nearest van der Waals protein contact is shown as a grey surface. Two subunits are depicted as ribbons; three are omitted for clarity. Amino acids in the neck and aperture regions are drawn as gray sticks; Ca^2+^ ions are cyan spheres. Approximate boundaries of the lipid membrane are indicated. **c**, Pore dimensions in the open and closed conformations. Dashed lines indicate the diameters of a dehydrated (black) and hydrated (cyan) Cl^−^ ion. **d**, Superposition of individual subunits from the closed and open conformations with a-helices depicted as cylinders. The boxed area shows a close-up of the neck region, with residues depicted as sticks.

In the closed conformation, the neck is less than 3.5 Å in diameter and approximately 15 Å long; three hydrophobic amino acids on the neck helix (S2b) from each of the channel’s five subunits, I76, F80 and F84, form its walls (Fig. 1a, c, d) (*6*). Its narrowness, length and hydrophobicity create an energetic barrier to ion permeation that seals the channel shut by hydrophobic block (*16*). In the open conformation, the neck has dilated to approximately 13 Å in diameter, which is more than sufficient to allow permeation of hydrated Cl^−^ ions (Fig. 1c). No appreciable conformational difference is present in the cytosolic region of the channel, and in particular, the aperture constriction of the pore retains its dimensions.

Comparison of the open and closed conformations of the neck highlights an unusual structural element of the pore that distinguishes the mechanism of gating in BEST1 from most other channels. The neck helix (S2b) is flanked on both of its ends by disruptions of α-helical secondary structure. These disruptions provide impressive flexibility on the one hand and tethering on the other that allow the helix to “float” between closed and open conformations (Fig. 2). In the closed conformation, hydrophobic packing at the center of the neck among the I76, F80 and F84 residues themselves stabilize this conformation. In the open conformation, the tendency of the phenylalanine residues to seclude their hydrophobicity from an aqueous environment is satisfied by their interactions with other hydrophobic amino acids (Y236 and W287) on the S3b and S4a helices located behind the neck helix (Fig. 2 b). The conformational change moves F80 and F84 away from the center of the pore and involves a slight rotation along the helical axis of S2b (~ 10 ° clockwise viewed from the extracellular side), outward displacement of S2b (~ 2.5 Å at F80), a slight expansion of the entire transmembrane region (~ 1 Å increase in radius), and a coordinated set of side chain rotamer changes (Fig. 2a, b and video). Both F80 and F84 move from the most commonly observed rotamer for phenylalanine (observed 44% of the time in the pdb) in the closed conformation to the second most commonly-observed rotamer conformation (observed 33% of the time) in the open conformation (*17*). By these conformational changes, F80 and F84 rotate away from the axis of the pore by 80° and 105°, respectively (Fig. 2, fig. S5). In a domino effect, side chain rotamer changes of Y236, F282, F283, and W287 allow for the movements of F80 and F84 (Fig. 2a, b and movie S1). The conformational change in I76 is also dramatic. When the channel opens, the first α-helical turn of the neck helix unravels such that I76 packs with F247, F276, L279 and F283 in the open conformation and has shifted by approximately 10 Å (fig. S5a, d). The unraveling is facilitated by P77, which is perfectly conserved among BEST channels and is part of the neck helix in its closed conformation, but marks its N-terminal end in the open conformation (Fig. 2c, d). The repositioning within the neck also exposes S79 and G83 on the neck helix, which are secluded behind the F80 and F84 in the closed conformation, to the pore in the open conformation (Fig. 2a, b and video). Thus, through a concertina of coordinated conformational changes in and around the neck helix, amino acids that formed the hydrophobic barrier that prevented ion permeation have dispersed and reveal a wide aqueous vestibule.

**Fig. 2.**
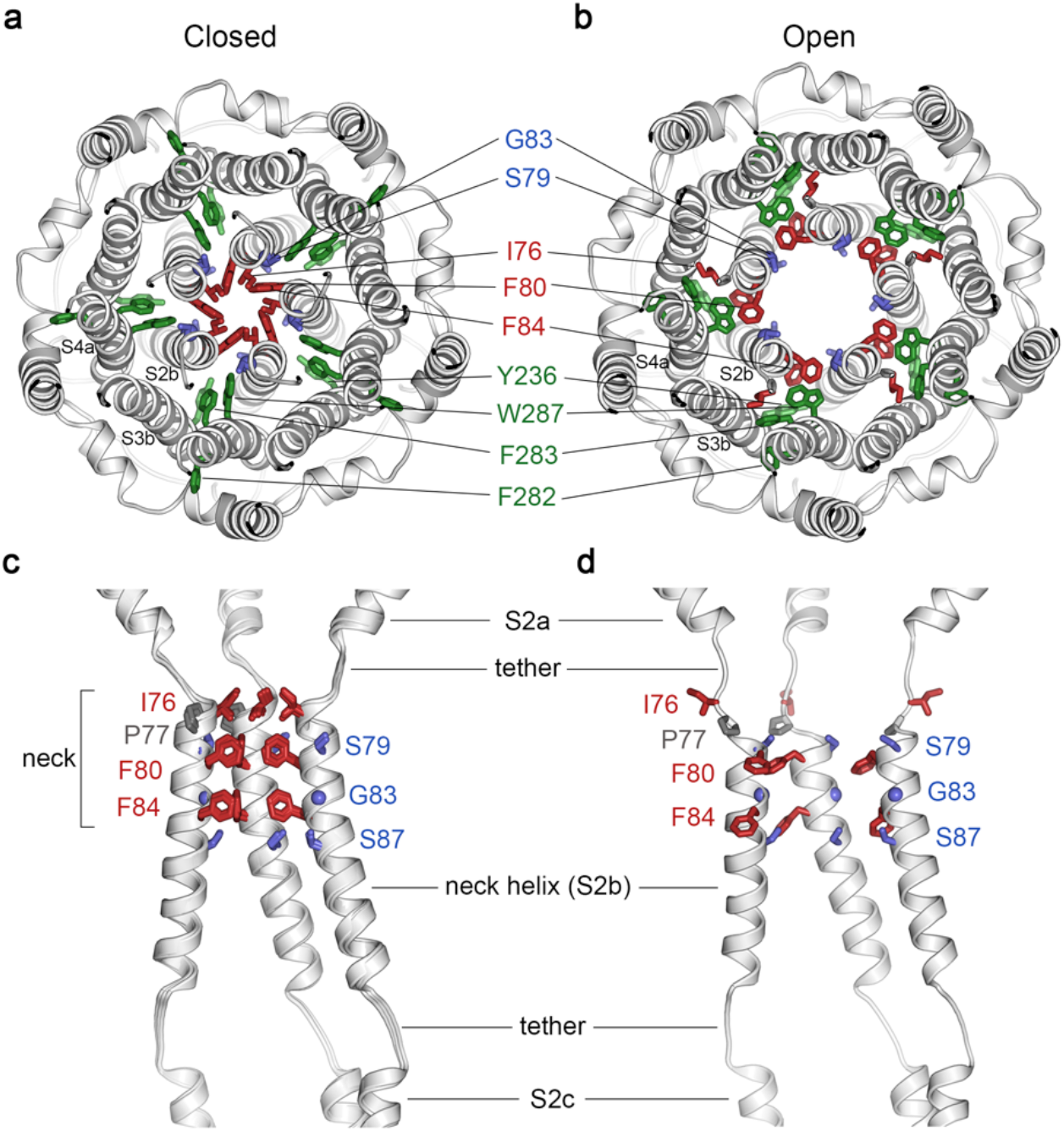
Opening transitions. **a-b**, Cutaway views of the neck region for the closed (a) and open (b) conformations, viewed from the extracellular side and shown as ribbons. Residues that form the hydrophobic seal in the closed conformation (I76, F80, F84) are colored red in both conformations. Surrounding aromatic residues that move to accommodate opening are colored green. Residues that become exposed to the pore in the open conformation (S79, sticks, and G83, sphere) are blue. A supplementary video shows the transition. **c-d**, Side view of the conformational changes in the neck; closed (c) and open (d). In (c), a superposition of the structures of BEST1_405_ in the Ca^2+^-bound inactivated conformation, BEST1_345_ in the Ca^2+^-bound closed conformation, and BEST1_345_ in the Ca^2+^-free closed conformation shows that the neck adopts an indistinguishable (closed) conformation in each. The S2a, b, and c helices from three subunits are shown. Residues are depicted and colored as in a-b; P77 is gray; S87 is shown for reference.

To address how Ca^2+^ binding contributes to BEST1 channel gating, we determined the cryo-EM structure of BEST1_345_ in the absence of Ca^2+^ to 3.0 Å resolution (fig. S6, fig. S7; a cryo-EM structure of BEST1_405_ without Ca^2+^ was also determined, at 3.6 Å resolution, and is indistinguishable). The Ca^2+^-free structure is very similar to the inactivated and Ca^2+^-bound closed conformations with the only notable differences near the Ca^2+^ clasp (fig. S8). Of particular note, the neck shares the same closed conformation as in the inactivated and Ca^2+^- bound closed structures (Fig. 2c). In structures with Ca^2+^ bound, the five Ca^2+^ clasps, one from each subunit, resemble a belt that wraps around the midsection of BEST1 (Fig. 1a, b). Without Ca^2+^, the majority of the Ca^2+^ clasp becomes disordered (fig. S8b). 3D classification of the Ca^2+^-free dataset yielded only closed conformations of the neck but did indicate a degree of flexibility in the channel between the transmembrane and cytosolic regions that was manifested as a ~ 5° rotation along the symmetry axis (fig. S8c). This conformational flexibility was not observed in the Ca^2+^-bound datasets, which suggests that Ca^2+^ binding rigidifies the channel and we hypothesize that this may be necessary for stabilization of the open conformation.

To investigate how the coupling of the amino acid side chain movements are involved in the transition between the open and closed conformations of the neck and how Ca^2+^ binding might bias these, we studied the effects of mutating W287 to phenylalanine. W287 is highly conserved among BEST channels. We chose to study this residue because it adopts one side chain rotamer and packs with both F80 and F84 in the open conformation of the neck and another side chain rotamer in the closed conformation that buttresses the space between adjacent S2b helices (Figs. 2a, b and 3d, e), and thus it might govern conformational changes in the neck. We find that the W287F mutation produces channels with dramatically altered gating. Whilst the W287F mutant retained normal Cl^−^ versus K^+^ selectivity (Fig. 3a), the mutation makes the channel nearly insensitive to Ca^2+^; approximately 80% of the Cl^−^ current level was maintained when Ca^2+^ was chelated with EGTA (Fig. 3a, b). To understand the molecular basis of this behavior, we determined cryo-EM structures of the W287F mutant in the presence and absence of Ca^2+^ at 2.7 Å and 3.0 Å resolutions, respectively (fig. S7, 9). In Ca^2+^, the channel adopts an open conformation that is essentially indistinguishable from the open conformation observed for BEST1_345_ (fig. S9, Cα RMSD = 0.2 Å). Unlike BEST1_345_, 3D classification did not reveal a closed conformation within the cryo-EM dataset, which indicates that essentially all of the particles are in an open conformation and that the open state is preferential for this mutant.

**Fig. 3.**
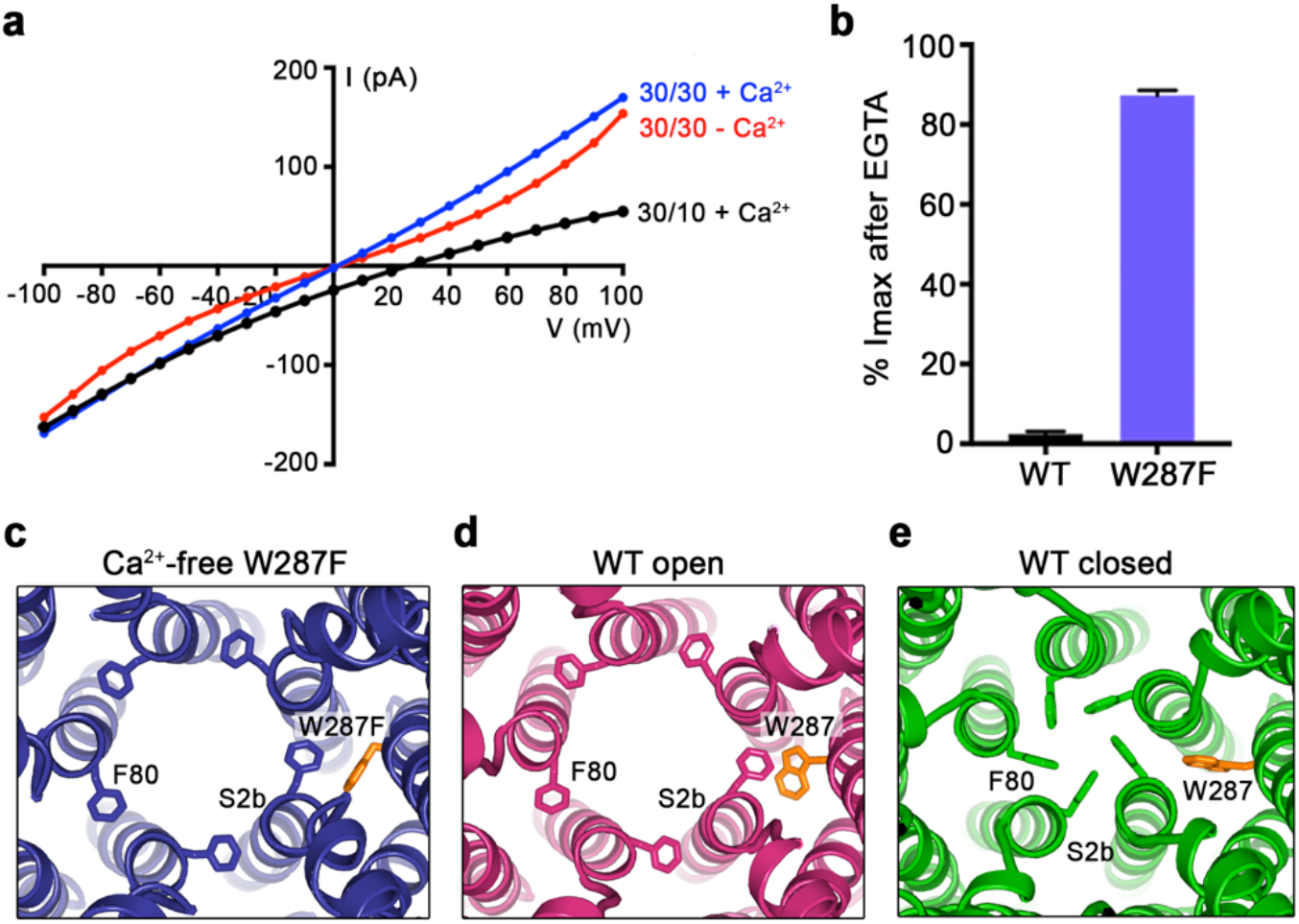
The W287F mutant decouples the Ca^2+^ ligand from the activation gate. **a-b**, Dramatically reduced Ca^2+^-dependence but normal Cl^−^ versus K^+^ selectivity of the W287F mutant. *I-V* relationships (a) are shown for voltages stepped from −100 to +100 mV for the indicated conditions [cis/trans KCl concentration in mM, and ~ 300 nM [Ca^2^]_free_ (+Ca^2+^) or 10 mM EGTA (- Ca^2+^)]. The reversal potential (E_rev_) measured using asymmetric KCl (30/10 mM) indicates normal Cl^−^ versus K^+^ selectivity: E_rev_ = 24.8 ± 0.8 mV for BEST1_345_ W287F in comparison to 23.4 ± 0.3 mV for BEST1_345_ [Vaisey and Long, *JGP, in press]*). **b**, Bar graph showing the percentage of current remaining after addition of 10 mM EGTA for BEST1_345_ (WT) and the W287F mutant. I_max_ indicates the current measured at +100 mV in the presence of 300 nM [Ca^2+^]_free_. Error bars denote the SEM calculated from four (WT) or six (W287F) separate experiments. **c-e**, The W287F mutant locks the neck open, even in the absence of Ca^2+^. c, Structure of the W287F mutant in the absence of Ca^2+^, showing the open conformation of the neck region (ribbons, cutaway view from an extracellular orientation). Density for the Ca^2+^ clasp is disordered indicating that Ca^2+^ is not bound in the structure (Extended Data Fig. 9). The W287F mutation (orange sticks) is shown for one subunit. F80 residues are drawn as sticks. Open (d) and closed (e) conformations of Ca^2+^-bound BEST1_345_ are depicted in the same manner.

In the absence of Ca^2+^, in spite of missing density for Ca^2+^ and a disordered Ca^2+^-clasp region, the neck also adopts an open conformation (Fig 3c and fig. S9). Thus, in accord with the electrophysiological recordings, the W287F mutation decouples Ca^2+^ binding from the conformational changes in the activation gate. Modeling of the W287F mutation on a closed conformation of the channel introduces a void behind the neck (fig. S9h), which we hypothesize energetically disfavors the closed conformation. The effects of the relatively conservative mutation of tryptophan to phenylalanine give context to a myriad of disease-causing mutations in and around the neck (fig. S5e).

The structures reveal that the open pore of BEST1 comprises a 90 Å-long water-filled vestibule and a short constriction at the cytosolic aperture (Fig. 1b). The aperture constriction is only 3 Å long (measured where the pore diameter is < 4 Å); its walls are formed solely by the side chains V205 of the five subunits (Fig. 4a). Retinitis pigmentosa can be caused by mutation of the corresponding residue of human BEST1 (I205T mutation)(*18*), which suggests that the aperture has an important role in channel function. The structures reveal that the aperture has the same conformation in the open and closed states (Fig. 1a, b); accordingly the V205A mutation of chicken BEST1, which would be expected to widen the aperture markedly, has no effect on Ca^2+^-dependent activation or inactivation (*7, 15*). We conclude that the aperture does not function as the activation or inactivation gate. However, mutations of V205 have dramatic effects on ion permeability(*7*) (Fig 4c, d). Both human and chicken BEST1 have a lyotropic permeability sequence in which small anions that are more easily dehydrated than Cl^−^, such as Br^−^, I^−^ and SCN^−^, are more permeable(*7, 19–21*) (Fig. 4c). We find that mutation of V205 to a smaller or more hydrophilic residue (e.g. glycine, alanine, or serine) abolishes the lyotropic sequence whereas mutation to isoleucine, a bulkier hydrophobic amino acid, makes the permeability differences between Cl^−^, Br^−^, I^−^ and SCN^−^ more dramatic (Fig. 4c). Thus, the aperture controls permeability among anions. Based on the narrow diameter of the aperture, anions would shed at least some of their water molecules as they pass through it; this would give rise to the channel’s lyotropic permeability sequence and may contribute to its low single channel conductance (reported at ~2 μS for Cl^−^ for drosophila BEST1(*22*)). The permeability to large anions such as acetate, propionate and butyrate increases when V205 is substituted by alanine or glycine (Fig. 4d), and thus, as has been suggested previously(*7*), the aperture functions as a size-selective filter that would tend to prevent permeation of large cellular constituents such as proteins or nucleic acids. Notably, the amino acid sequence at and around the aperture varies among BEST channels (Fig. 4b), and this may endow these channels with distinct permeabilities related to their specific physiological functions. Data suggest that the inhibitory neurotransmitter GABA permeates through BEST1 to underlie a tonic form of synaptic inhibition in glia (*13*). While this possibility seemed incongruous with the narrowness of the neck observed in the initial structure, the widened neck of the open conformation and the presence of a single constriction that controls permeability make the possibility of slow conductance of GABA and/or other solutes of similar size more plausible. Although the aperture adopts an indistinguishable conformation in all of the structures, we suspect that “breathing” (e.g. thermal motions) of the protein could allow larger ions to move through the aperture than might otherwise fit. It is also conceivable that the binding of cellular ligands near the aperture, as has been suggested for ATP (*23*), could influence channel behavior by changing its dimensions somewhat. The structure of the open pore hints at a rich diversity of potential physiological functions for BEST channels that are largely unexplored.

**Fig. 4.**
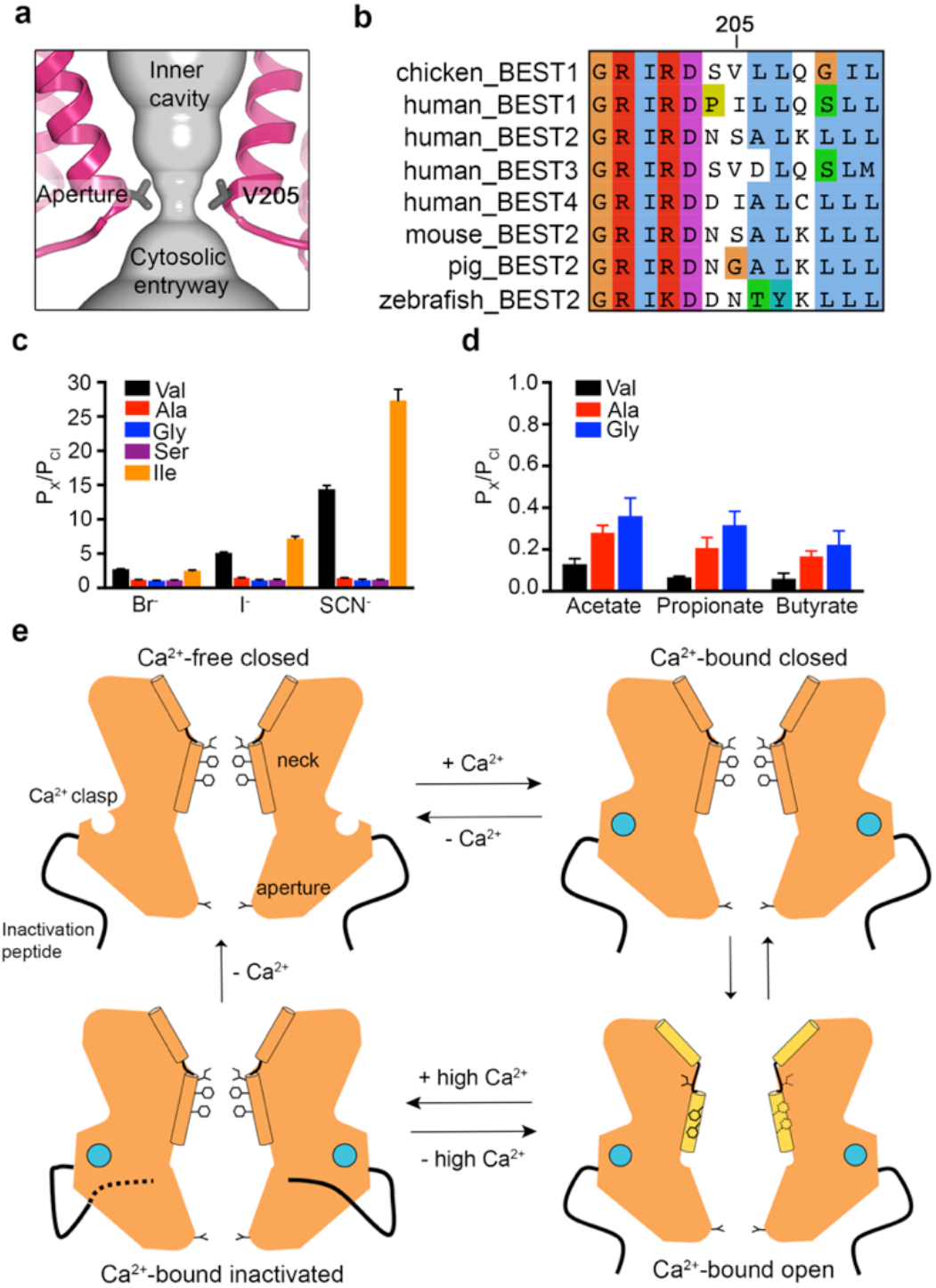
The aperture and a gating model. **a**, Close up of the aperture. **b**, Sequence alignment around the aperture. **c-d**, Mutation of V205 affects ion permeability. c, Comparison of the permeabilities of Br^−^, I^−^, and SCN^−^ relative to Cl^−^ (P_x_/P_cl_) for wild type (Val) and the indicated mutants of V205. P_x_/P_cl_ values were calculated from reversal potentials recorded in 30 mM KCl (cis) and 30 mM KX (trans) where X is Br, I, or SCN. IV traces are shown in Fig. S10. d, Permeabilities of larger anions. Comparison of the permeabilities of acetate, propionate and butyrate relative to Cl^−^ for wild type (Val) and the indicated mutants of V205 (calculated as in c). For c-d, error bars denote the SEM from three experiments. **e**, Gating model. In the absence of Ca^2+^, hydrophobic block at the neck prevents ion flow (Ca^2+^-free closed). When the Ca^2+^ clasps are occupied by Ca^2+^, the channel is in equilibrium between Ca^2+^-bound closed and Ca^2+^-bound open conformations. The dramatic widening of the opened neck enables hydrated ions to flow through it. Binding of the inactivation peptide to its cytosolic receptor, which is stimulated by higher concentrations (> 500 nM) of Ca^2+^, induces the Ca^2+^-bound inactivated conformation in which the neck is closed. The aperture, which remains fixed throughout the gating cycle, acts as a size-selective filter that requires permeating ions to become partially dehydrated as they pass though it, and this engenders the channel’s lyotropic permeability sequence.

The structures presented herein represent the major gating transitions in the channel (Fig. 4e). Unlike voltage-dependent K^+^ and Na^+^ channels, in which ions are prevented from flowing by different mechanisms in the inactivated and deactivated states (*21, 24*), the same closed conformation of the neck is responsible for the deactivated (Ca^2+^-free) and inactivated states of BEST1. While localized twisting or domain motions often constitute the activation mechanism of ion channels, dramatic molecular choreography within the protein core of BEST1 underlies opening and represents a new paradigm for ion channel gating.

## Materials and Methods

### Cloning, expression and purification of BEST1

Chicken BEST1 (UniProt E1C3A0) constructs (amino acids 1–405 or 1–345 followed by a Glu-Gly-Glu-Glu-Phe tag) were expressed in *Pichia Pastoris* as described previously(*6*). Mutations were made using standard molecular biology techniques.

In preparation for cryo-EM analysis, purification of BEST1 proteins were performed as described previously with modification (*6*). BEST1 protein was purified by size-exclusion chromatography (SEC; Superose 6 increase 10/300 GL; GE Healthcare) in buffer containing 20 mM Tris, pH 7.5, 50 mM NaCl, 1 mM *n*-dodecyl-β-D-maltopyranoside (DDM; Anatrace) and 0.1 mM cholesteryl hemisuccinate (CHS; Anatrace). Purified BEST1 was concentrated to 5 mg ml^−1^ using a 100 kDa concentrator (Amicon Ultra-4, Millipore) and divided into aliquots. For structures with Ca^2+^, 1 μM CaCl_2_ was added to the freshly purified protein. For Ca^2+^-free structures, and 5 mM EGTA, pH 7.5 was added to the freshly purified protein. These samples were immediately used for cryo-EM grid preparation.

### EM sample preparation and data acquisition

5 μl of sample was pipetted onto Quantifoil R1.2/R1.3 holy carbon grids (Au 400, Electron Microscopy Sciences), which had been glow discharged for 10 s using a PELCO easiGlow glow discharge cleaning system (Ted Pella). A vitrobot Mark IV cryo-EM sample plunger (FEI) (operated at room temperature with a 1–2 s blotting time under a blot force of 0 and 100% humidity) was used to plunge-freeze the sample into liquid nitrogen-cooled liquid ethane. For Ca^2+^-free conditions, the blotting paper used for grid freezing was pre-treated with 2 mM EGTA solution (4x), rinsed with ddH_2_O (4x) and dried under vacuum. Grids were clipped and loaded into a 300 keV Titan Krios microscope (FEI) equipped with a K2 Summit direct electron detector (Gatan). Images were recorded with SerialEM(*25*) in super-resolution mode at a magnification of 22,500x, which corresponds to a super-resolution pixel size of 0.544 Å, and a defocus range of −0.7 to −2.15 μm. The dose rate was 9 electrons per physical pixel per second, and images were recorded for 10 seconds with 0.25 s subframes (40 total frames), corresponding to a total dose of 76 electrons per Å^2^.

### Image processing

Figures S1, S2, S6 and S9 show the cryo-EM workflow for Ca^2+^-bound BEST1_405_, Ca^2+^-bound BEST1_345_, Ca^2+^-free BEST1_345_ and BEST1_345_ W287F with and without Ca^2+^, respectively. Movie stacks were gain-corrected, two-fold Fourier cropped to a calibrated pixel size of 1.088 Å, motion corrected and dose weighted using MotionCor2(*26*). Contrast Transfer Function (CTF) estimates for motion-corrected micrographs were performed in CTFFIND4 using all frames (*27*).

#### *Ca^2+^-bound BEST1*_405_, *BEST1*_345_ *and BEST1*_345_ *W287F datasets*

All subsequent image processing was carried out with RELION2.1(*28*), using a particle box size of 384 pixels and a spherical mask with a diameter of 140–160 Å. A total of 1740, 1644 and 1597 micrographs were collected for Ca^2+^-bound BEST1_405_, BEST1_345_ and BEST1_345_ W287F, respectively, and all micrographs were inspected manually; poor quality micrographs and those having CTF estimation fits lower than 5 Å were discarded. Approximately 1000 particles were selected manually for reference-free 2D classification to generate templates that were then used for automatic particle picking. Auto-picking yielded ~312,000, ~309,000 and ~308,000 particles for BEST1_405_, BEST1_345_ and BEST1_345_ W287F, respectively. One round of 2D classification, using 100 classes, was used to remove outlier particles (e.g. ice contaminants), and this yielded ~290,000 particles for BEST1_405_ and BEST1_345_ datasets and ~265,000 particles for BEST1_345_ W287F. 3D refinement, using C5 symmetry, was performed for each dataset using an initial model (generated from a previously collected, lower resolution cryo-EM dataset of Ca^2+^-free BEST1 using EMAN2(*29*)) that was low-pass filtered at 60 Å resolution. This yielded consensus reconstructions at 3.1 Å (BEST1_405_)and 2.9 Å (BEST1_345_) overall resolutions that have the closed conformation of the neck and 2.8 Å for BEST1_345_ W287F that an open conformation at the neck. (Refinement using C1 symmetry also yielded reconstructions with 5fold symmetry.) All overall resolution estimates are based on gold-standard Fourier shell correlations (FSC).

To identify the distinct conformational states within the Ca^2+^-bound BEST1_345_ dataset, we performed 3D classification using the consensus reconstruction as an initial model (low-pass filtered at 5 Å resolution) and sorting the particles into 9 classes. One class with a widened neck (BEST1_345_ open) was isolated, containing ~ 30,000 particles. To identify additional open particles from the dataset, this reconstruction was low-pass filtered at 5 Å resolution and used as an initial model for 3D classification (with 4 classes) on the entire dataset. This procedure yielded one class in the open conformation (containing approximately 40,000 particles) and three classes in the Ca^2+^-bound closed conformation (containing the remainder of the particles). One class for the closed conformation was chosen (containing approximately 44,000 particles) because it contained better-resolved density for the residues lining the neck (I76, F80, F84). 3D Refinement of these two classes yielded reconstructions at 3.0 Å overall resolution. Particles from these two classes were “polished” using aligned movie frames generated from MotionCor2(*26*). 3D refinement using the polished particles and a global angular sampling threshold of 1.75° yielded final reconstructions at 3.0 Å and 2.9 Å overall resolutions for the Ca^2+^-bound closed and open reconstructions of BEST1_345_, respectively. The same polishing strategy for the BEST1_345_ W287F dataset yielded a final reconstruction of 2.7 Å. Several analogous 3D classification procedures were performed to try to identify an open conformation in the Ca^2+^-bound BEST1_405_ dataset but none were found. Conversely, 3D classification approaches with the Ca^2+^-bound BEST1w287f dataset to identify multiple conformations yielded only reconstructions with an open neck.

#### *Ca^2+^-free BEST1*_345_ *and* Ca^2+^*-free BEST1*_345_ *W287F datasets*

Initial image processing was carried out with RELION2.1(*28*), using a particle box size of 384 pixels and a mask diameter of 140. A total of ~1000 or ~1600 micrographs were collected for Ca^2+^-free BEST1_345_ and Ca^2+^-free BEST1_345_ W287F, respectively, and manually pruned as described for the Ca^2+^-bound dataset. Auto-picking templates were generated as described and the selected particles (~150,000 for the Ca^2+^-free BEST1_345_ and ~185,000 particles for the BEST1_345_ W287F datasets) were subjected to one round of 2D classification with 100 classes. 3D refinement was performed using the selected particles from 2D classification (~130,000 for the Ca^2+^-free BEST1_345_ and ~150,000 particles for the Ca^2+^-free BEST1_345_ W287F datasets), C5 symmetry, and the EMAN2-generated initial model. This yielded reconstructions of 3.4 Å and 3.2 Å overall resolutions, respectively. Particle polishing was performed on each dataset and the polished particles were imported into the cisTEM cryo-EM software package for further refinement and classificatio(*30*) 3D refinement was performed in cisTEM using a mask to apply a 15 Å low-pass filter to the micelle region. This yielded final reconstructions to 3.0 Å overall resolution for both datasets. The ~ 5 ° relative rotation of the cytosolic region with respect to the transmembrane region that was observed under Ca^2+^-free conditions was identified using 3D classification (using 6 or 8 classes for Ca^2+^-free BEST1_345_ and Ca^2+^-free BEST1_345_ W287F, respectively) using spatial frequencies up to 6 Å for refinement. Refinement of 3D classes with the most extreme rotation (e.g. approximately ± 2.5 ° rotations relative to the consensus reconstruction) in cisTEM yielded overall resolutions of 3.6 Å (Ca^2+^-free BEST1_345_ conformation A, ~11,000 particles), 3.4 Å (Ca^2+^-free BEST1_345_ conformation B, ~21,000 particles), 3.4 Å (Ca^2+^-free BEST1_345_ W287F conformation A, ~21,000 particles) and 3.5 Å (Ca^2+^- free BEST1_345_ W287F conformation B, ~17,000 particles). These reconstructions for Ca^2+^-free BEST1_345_ are depicted in Supplementary Figure 8.

RELION2.1(*28*) was used to estimate of the local resolution all of the final maps. The maps shown in figures are combined maps, were sharpened (using a 5-factor of −50–75 Å^2^), and low-pass filtered at the final overall resolution of each map.

### Model building and refinement

The atomic models were manually built into one of the half-maps (which had been sharpened using a 5-factor of −50–75 Å^2^ and low-pass filtered at the final overall resolution) using the X-ray structure of BEST1 as a starting point (PDB ID: 4RDQ) and were refined in real space using the COOT software(*31*). The atomic models were further refined in real space against the same half-map using PHENIX (*32*). The final models have good stereochemistry and good Fourier shell correlation with the other half-map as well as the combined map (Supplementary Figures 3 and 7). Structural figures were prepared with Pymol (pymol.org), Chimera (*33*), and HOLE (*34*).

### Liposome reconstitution

SEC-purified protein [in SEC buffer: 150 mM NaCl, 20 mM Tris-HCl, pH7.5, 3 mM *n*-decyl-β-D-maltoside (DM; Anatrace)] was reconstituted into liposomes. A 3:1 (wt/wt) mixture of POPE (Avanti) and POPG (Avanti) lipids was prepared at 20 mg ml^−1^ in reconstitution buffer (10 mM Hepes-NaOH, pH 7.6, 450 mM NaCl, 0.2 mM EGTA, 0.19 mM CaCl_2_). 8% (wt/vol) *n*-octyl-β-D-maltopyranoside (Anatrace) was added to solubilize the lipids and the mixture was incubated with rotation for 30 min at room temperature. Purified protein was mixed with an equal volume of the solubilized lipids to give a final protein concentration of 0.2–1 mg ml^−1^ and a lipid concentration of 10 mg ml^−1^. Proteoliposomes were formed by dialysis (using a 8000 Da molecular mass cutoff) for 1–2 days at 4 °C against 2–4 L of reconstitution buffer and were flash frozen in liquid nitrogen and stored at −80 °C until use.

### Electrophysiological recordings

Proteoliposomes were thawed and sonicated for approximately 10 s using an Ultrasonic Cleaner (Laboratory Supplies Company). All data are from recordings made using the Warner planar lipid bilayer workstation (Warner Instruments). Two aqueous chambers (4 mL) were filled with bath solutions. Chlorided silver (Ag/AgCl) wires were used as electrodes, submerged in 3 M KCl, and connected to the bath solutions via agar-KCl salt bridges [2% (wt/vol) agar, 3 M KCl]. The bath solutions were separated by a polystyrene partition with a ~200μM hole across which a bilayer was painted using POPE:POPG in *n*-decane [3:1 (wt/wt) ratio at 20 mg ml^−1^]. Proteoliposomes were applied to the bilayer with an osmotic gradient across the bilayer with solutions consisting of: 30 mM KCl or NaCl (*cis* side) and 10 mM KCl or NaCl (*trans* side), 20 mM Hepes-NaOH, pH 7.6 and 0.21 mM EGTA/0.19 mM CaCl_2_ ([Ca^2+^]_free_ ~ 300 nM) or 1 μM CaCl_2_. Proteoliposomes were added, 1 μL at a time, to the *cis* chamber to a preformed bilayer until ionic currents were observed. Solutions were stirred using a stir plate (Warner Instruments stir plate) to aid vesicle fusion. After fusion, the solutions were made symmetric by adding 3M KCl or 5M NaCl, depending on the starting solutions, to the *trans* side. Unless noted, all reagents were purchased from Sigma-Aldrich. All electrical recordings were taken at room temperature (22–24°C).

Measurements of relative permeabilities among anions were performed as described previously(*7*). Briefly, after establishing symmetric (30/30 mM KCl or NaCl) conditions, the bath solution in the *trans* chamber was replaced by perfusion with solutions in which KCl or NaCl was replaced by various potassium salts (Br, I, SCN, acetate, propionate) or sodium salts (butyrate).

Currents were recorded using the Clampex 10.4 program (Axon Instruments) with an Axopatch 200B amplifier (Axon Instruments) and were sampled at 200 μs and filtered at 1 kHz. Data were analyzed using Clampfit 10.4 (Axon Instruments). Graphical display and statistical analyses were carried out using GraphPad Prism 6.0 software. In all cases, currents from bilayers without channels were subtracted. Error bars represent the SEM of at least three separate experiments, each in a separate bilayer. We define the side to which the vesicles are added as the *cis* side and the opposite *trans* side as electrical ground, so that transmembrane voltage is reported as V_cis_-V_trans_. Ion channels are inserted in both orientations in the bilayer.

## Acknowledgements

We thank N. Grigorieff, members of his laboratory, and the staff at the Howard Hughes Medical Institute Cryo-EM facility for training and initial advice on cryo-EM. We thank M.J. de la Cruz of the Memorial Sloan Kettering Cancer Center Cryo-EM facility, M. Ebrahim, and the staff of the New York Structural Biology Center Simons Electron Microscopy Center for help with data collection. G.V received funding and mentorship from the Boehringer Ingelheim Fonds Predoctoral Fellowship Program.

## Funding

This work was supported, in part, by NIH Grant R01 GM110396 (to S.B.L) and a core facilities support grant to Memorial Sloan Kettering Cancer Center (P30 CA008748).

## Author contributions

A.N.M, G.V and S.B.L conceived of and designed the project. A.N.M determined structures of BEST1_405_, the W287F mutant, and structures in the absence of Ca^2+^. G.V determined structures of the Ca^2+^-bound closed state and the Ca^2+^-bound open state. A.N.M. and G.V performed electrophysiology experiments. All authors contributed to data analysis and the preparation of the manuscript.

## Competing interests

The authors declare no competing financial interests.

## Data and materials availability

Atomic coordinates and cryo-EM density maps of have been deposited with the PDB and Electron Microscopy Data Bank with the deposition numbers: D_1000236792 (BEST1_405_, inactivated), D_1000236798 (W287F mutant, Ca^2+^-free), D_1000236800 (W287F mutant, Ca^2+^-bound), D_1000236801 (Ca^2+^-free closed state), D_1000236802 (Ca^2+^-bound closed state), and D_1000236804 (Ca^2+^-bound open state). Correspondence and requests for materials should be addressed to S.B.L.

## Supplementary Materials

Materials and Methods

Figs. S1 to S10

References (25–34)

**Fig. S1.**
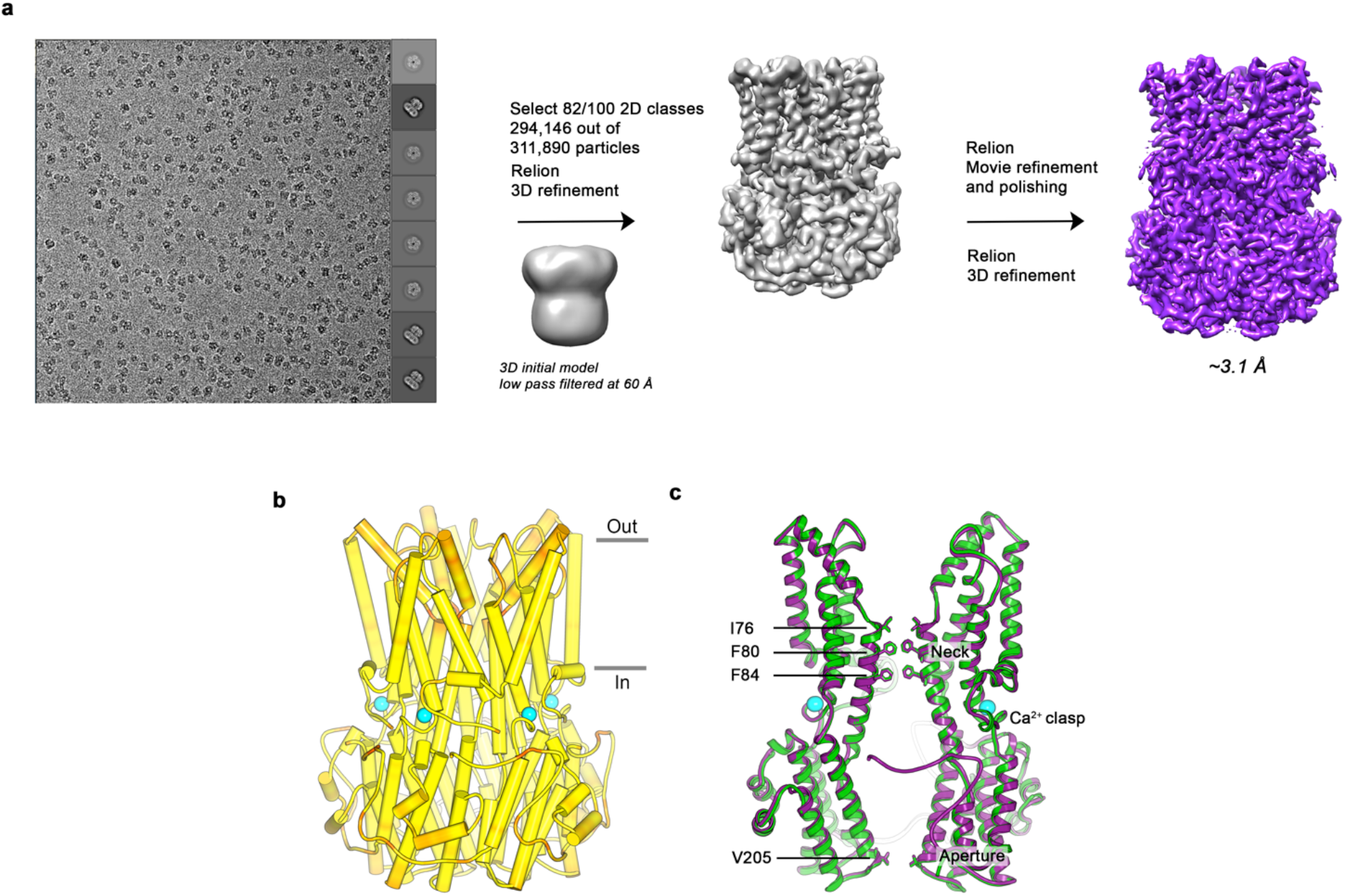
Cryo-EM workflow for the BEST1_405_ Ca^2+^-bound dataset and comparison of EM and X-ray structures. **a**, Cryo-EM workflow for the Ca^2+^-bound BEST1_405_ dataset (inactivated conformation). A detailed description can be found in the Methods. **b**, The X-ray and EM structures of BEST1_405_ are essentially indistinguishable. The structure of BEST1_405_ is drawn with a-helices depicted as cylinders and is colored on a yellow-to-red spectrum according to the displacement of Cα atoms between the BEST1_405_ cryo-EM structure and the X-ray structure (PDB ID: 4RDQ). Yellow represents displacements less than 0.5 Å and red represents displacements greater than 2 A. Ca^2+^ ions are depicted as cyan spheres and the approximate boundaries of a lipid membrane are indicated. **c**, A superposition shows that the Ca^2+^-bound closed conformation of BEST1_345_ (green) has the same overall conformation as BEST1_405_ (purple). Two subunits in ribbon representation are shown from the side. Ca^2+^ ions are drawn as cyan spheres and the labeled residues are shown as sticks.

**Fig. S2.**
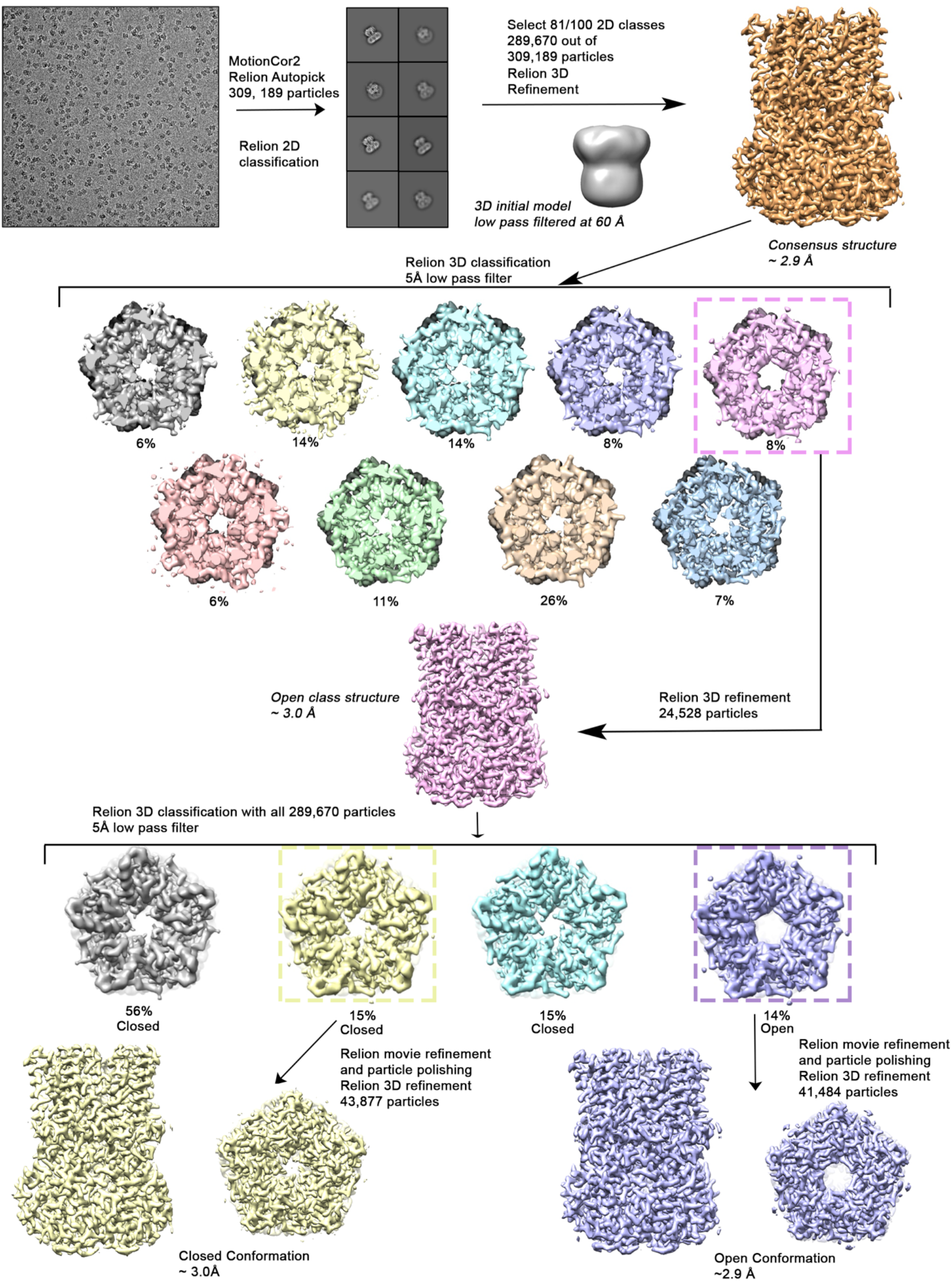
Cryo-EM workflow for the BEST1_345_ Ca^2+^-bound dataset. A detailed description can be found in Methods.

**Fig. S3.**
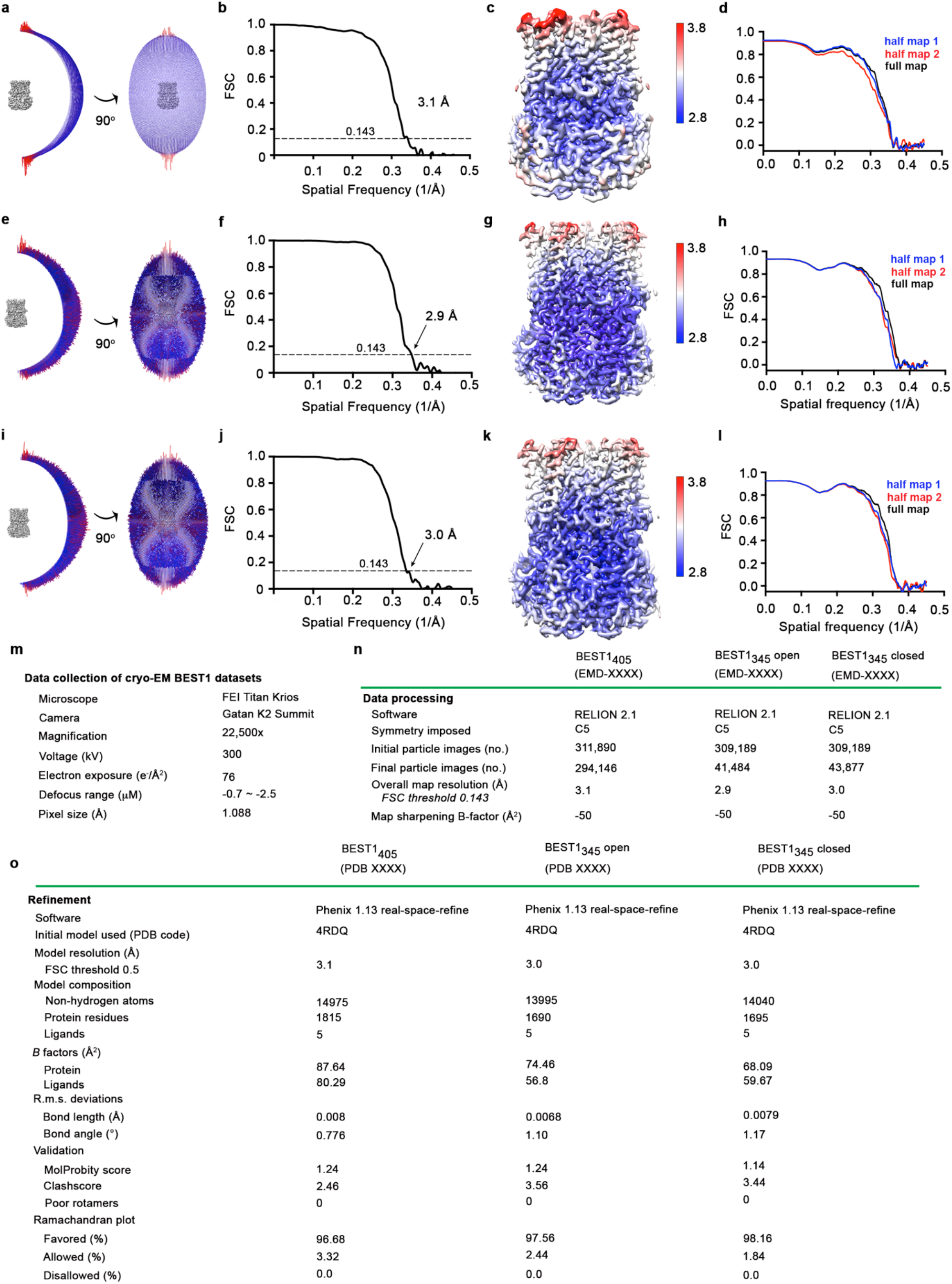
Structure determination of: Ca^2+^-bound BEST1_405_ (inactivated), Ca^2+^-bound open BEST1_345_, and Ca^2+^-bound closed BEST1_345_. **a-d**, Structural determination of BEST1_405_. **a**, Angular orientation distribution of particles used in final reconstruction. The particle distribution is indicated by color shading, with blue to red representing low and high numbers of particles. **b**, Gold-standard Fourier shell correlation (FSC) curve of the final 3D reconstruction. The resolution is 3.1 Å at the FSC cutoff of 0.143 (dotted line). **c**, Local resolution of the map was estimated using Relion^1^ and is colored as indicated. **d**, Model validation. Comparison of the FSC curves between the model and half map 1 (work), model and half map 2 (free) and model and full map. **e-h**, Structural determination of the Ca^2+^-bound open BEST1_345_ structure **e**, Angular orientation distribution of particles used in final reconstruction, similar to (a). **f**, Gold-standard Fourier shell correlation (FSC) curve of the final 3D reconstruction. The resolution is 2.9 Å at the FSC cutoff of 0.143 (dotted line). **g**, Local resolution of the map, as for (c). **h**, Model validation, as for (d). **i-l**, Structural determination of the Ca^2+^-bound closed BEST1 structure. **i**, Angular orientation distribution of particles used in final reconstruction, similar to (a). **j**, Gold-standard Fourier shell correlation (FSC) curve of the final 3D reconstruction. The resolution is 3.0 Å at the FSC cutoff of 0.143. **k**, Local resolution of the map, as in (c). **l**, Model validation, as in (d). **m-o**, Table of data collection and model statistics for the Ca^2+^-bound BEST1_405_ (inactivated), Ca^2+^- bound open BEST1_345_, and Ca^2+^-bound closed BEST1_345_ structures.

**Fig. S4.**
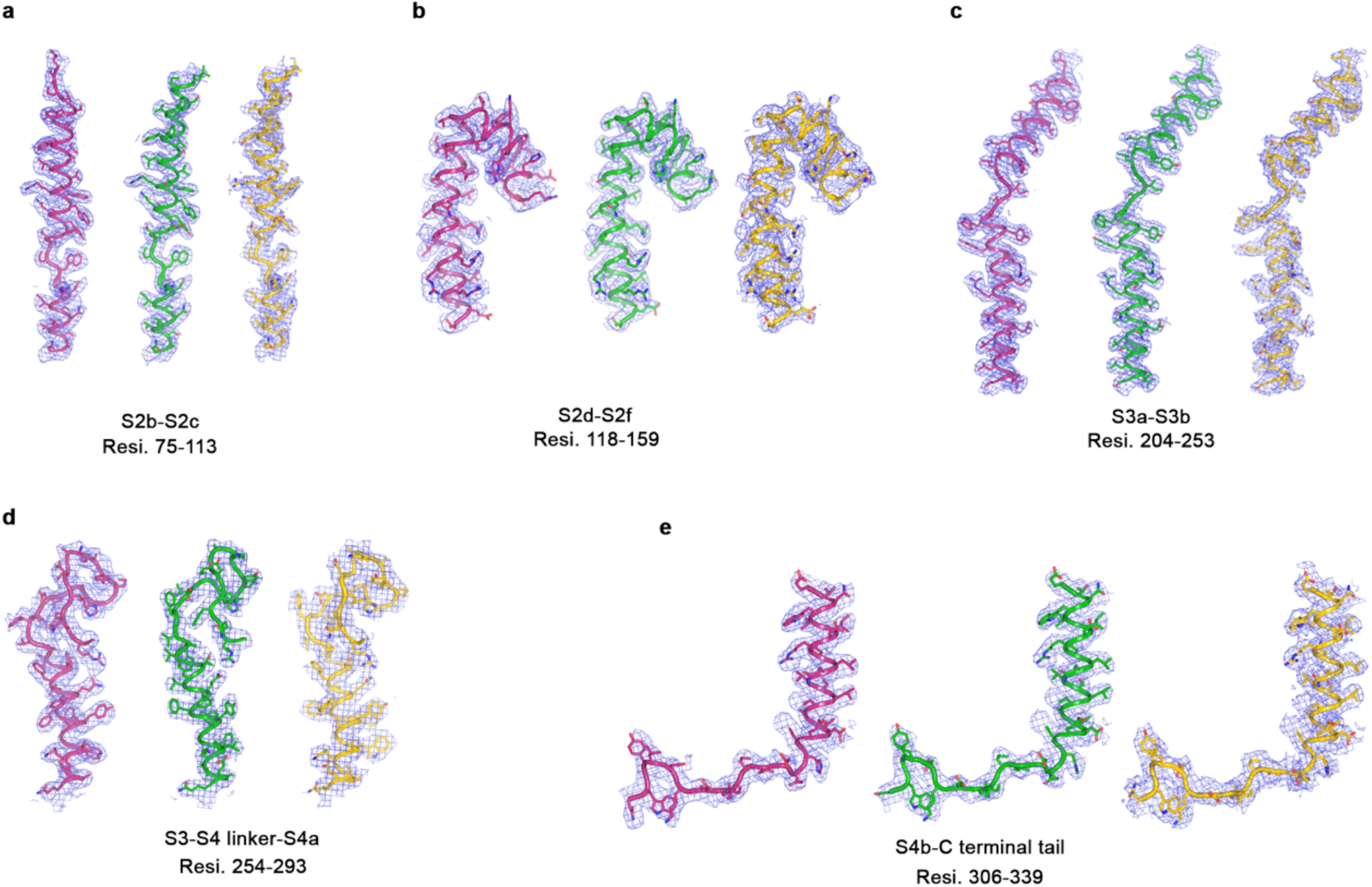
Representative cryo-EM density for three BEST1 cryo-EM structures. **a-e**, Representative map density (blue mesh, 5σ) highlighting different regions of channel in the Ca^2+^- bound open BEST1_345_ (pink), Ca^2+^-bound closed BEST1_345_ (green) and Ca^2+^-free BEST1_345_ (yellow).

**Fig. S5.**
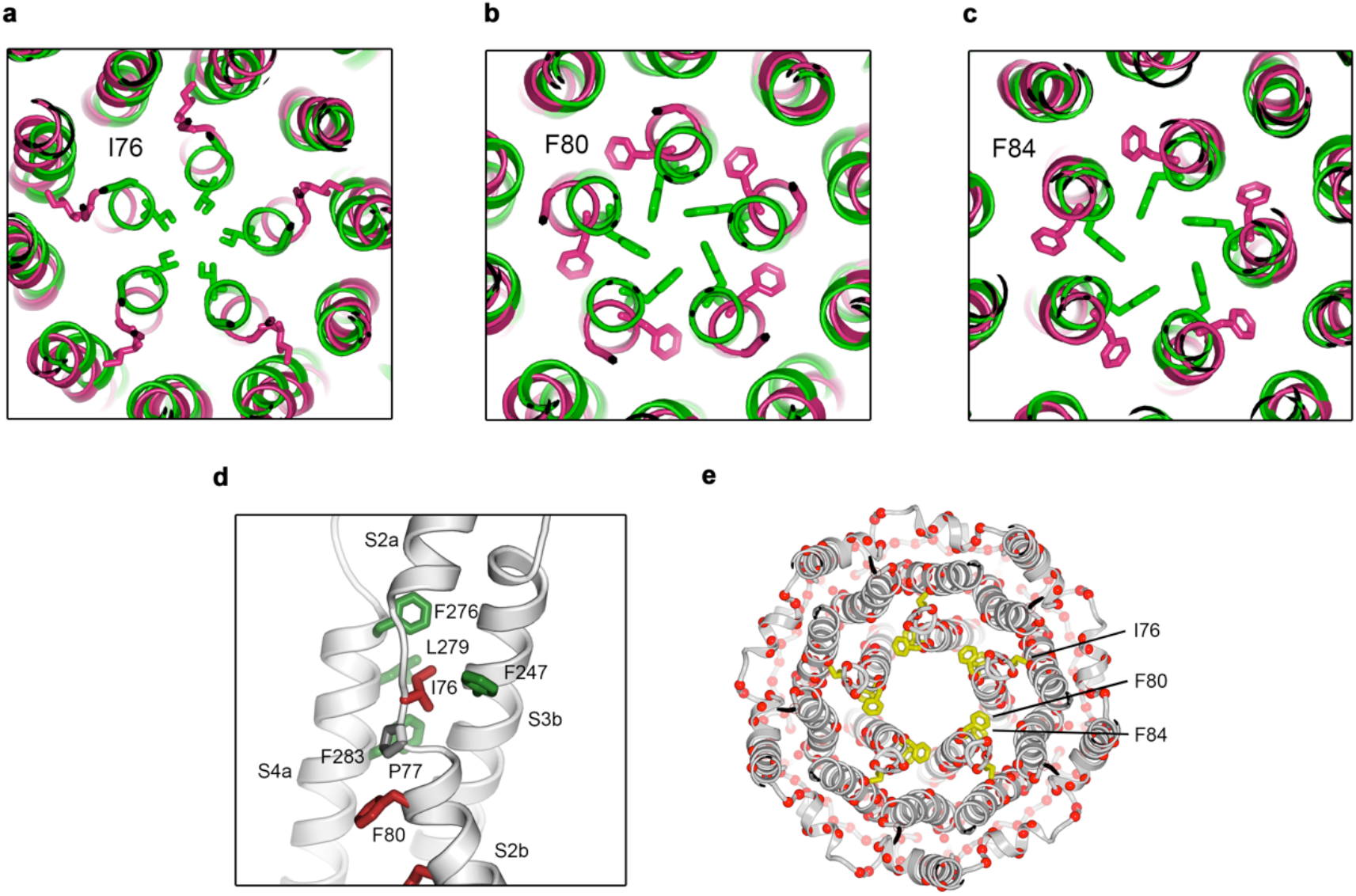
**a-c**, Comparison of neck-lining residues I76, F80 and F84 between the Ca^2+^-bound open (pink) and Ca^2+^-bound closed (green) structures. Side chains of labeled residues are depicted in each panel, viewed as a cutaway from the extracellular space. **d**, A close-up view showing the hydrophobic packing of I76 in the open conformation. Neck residues are highlighted in red, neighboring hydrophobic residues that interact with I76 are shown in green, and P77 is depicted in gray. **e**, Location of missense mutations associated with retinal diseases^31^ at amino acid positions in and around the neck of BEST1 (red spheres indicate the Cα positions of the mutations).

**Fig. S6.**
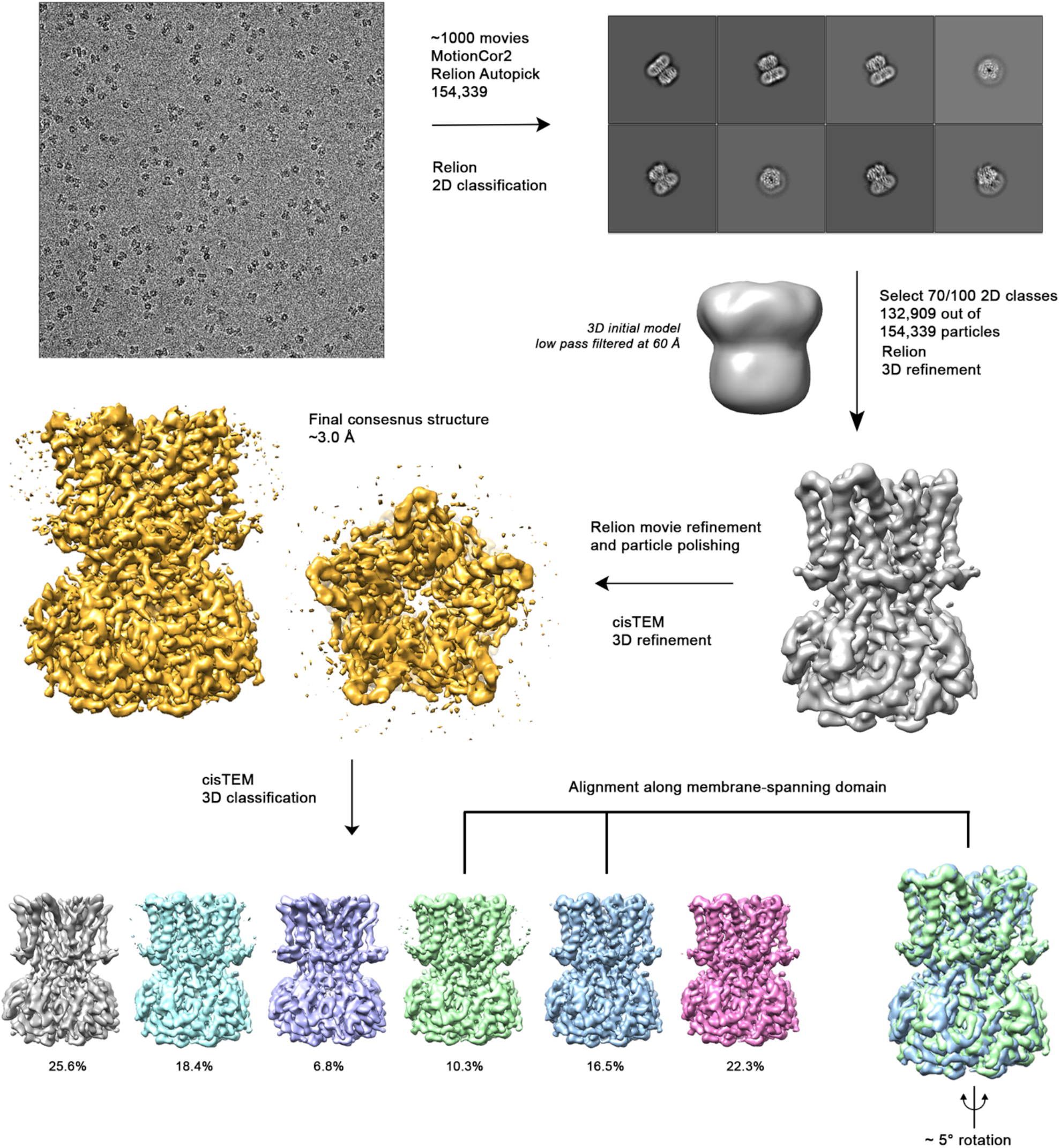
Cryo-EM workflow for the BEST1_345_ Ca^2+^-free dataset. A detailed description can be found in Methods.

**Fig. S7.**
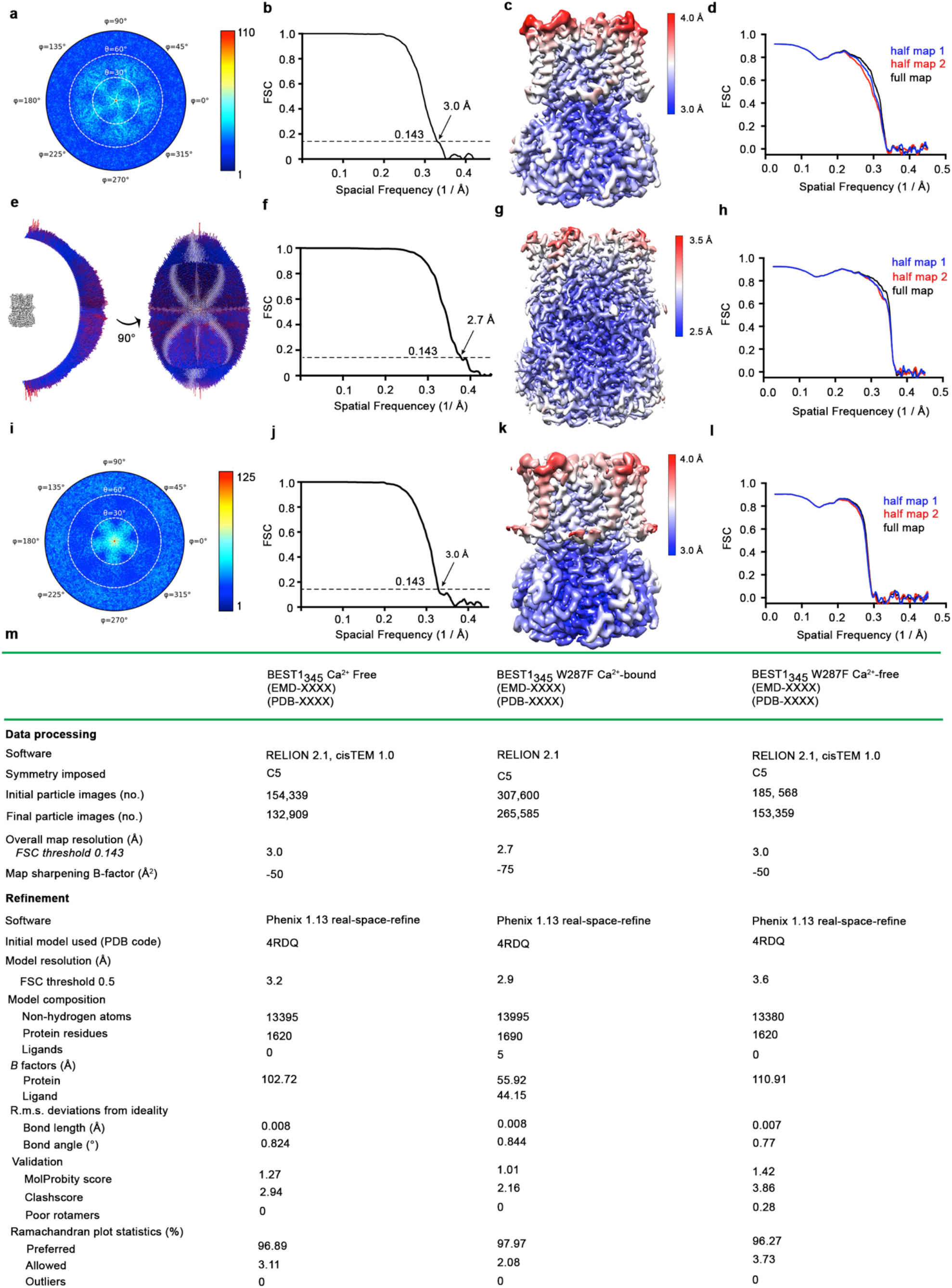
Cryo-EM structure determination of: Ca^2+^-free BEST1_345_, Ca^2+^-bound BEST1_345_ W287F, and Ca^2+^-free BEST1_345_ W287F. **a-d**, Structure determination of the consensus Ca^2+^- free BEST1_345_ conformation. **a**, Angular orientation distribution of particles used in the final reconstruction. The particle distribution is indicated by color shading, with blue to red representing low and high numbers of particles. **b**, Gold-standard Fourier shell correlation (FSC) curve of the final 3D reconstruction. The resolution is 3.0 Å at the FSC cutoff of 0.143 (dotted line). A thin vertical line indicates that only special frequencies to 1/(5 Å) were used to determine particle alignment parameters during refinement. **c**, Local resolution of the map estimated using Relion and colored as indicated. **d**, Model validation. Comparison of the FSC curves between the model and half map 1 (work), model and half map 2 (free) and model and full map are plotted. **e-h** Structure determination of the Ca^2+^-bound BEST1_345_ W287F structure as in *a-d.* **i-l**, Structure determination of the Ca^2+^-free BEST1_345_ W287F structure as in *a-d.* **m**, Table of data processing and model statistics.

**Fig. S8.**
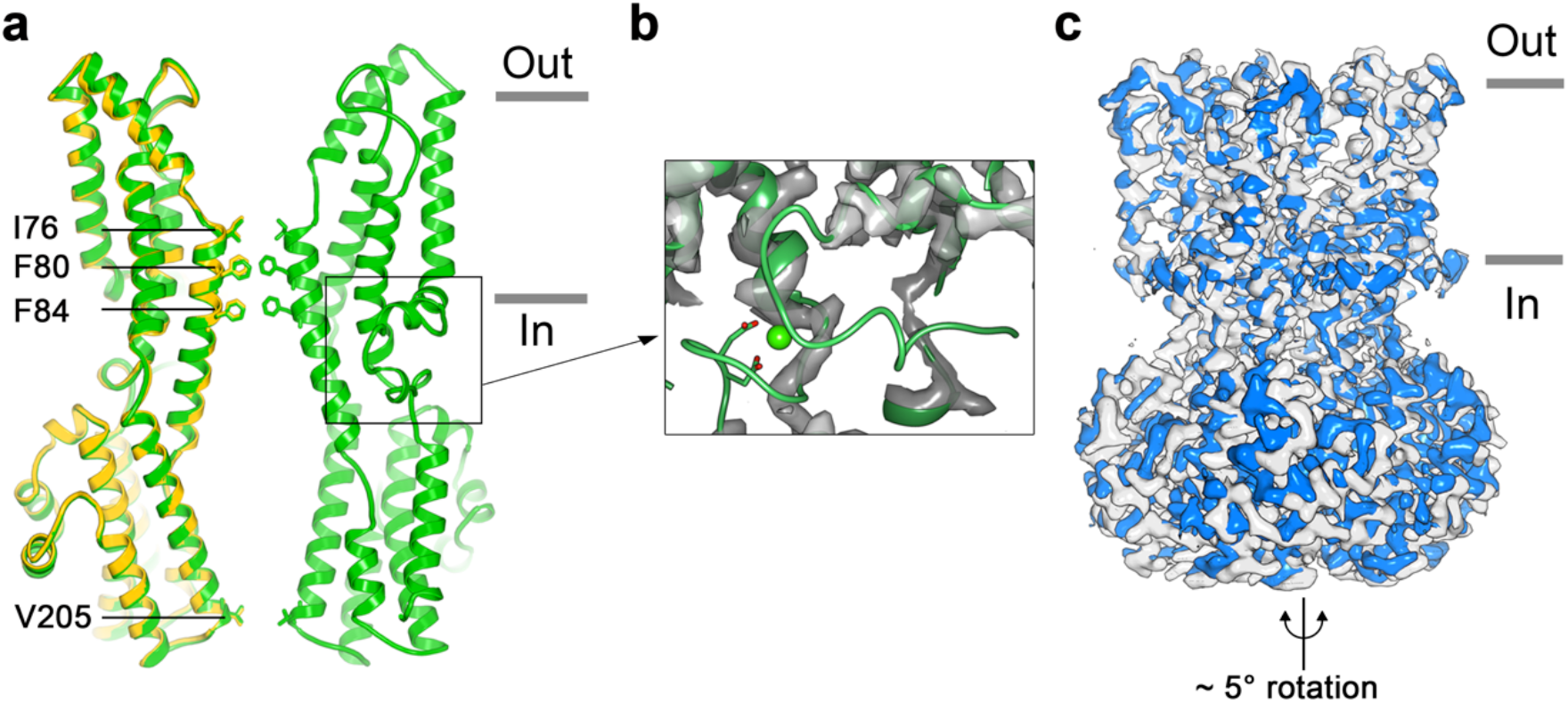
Structure of Ca^2+^-free BEST1_345_. **a**, Overlay comparison of the Ca^2+^-free conformation of BEST1_345_ (yellow) with the Ca^2+^-bound closed conformation of BEST1_345_ (green). One (Ca^2+^-free) or two (Ca^2+^-bound) channel subunits in ribbon are shown as a cutaway from the side with the approximate boundaries of the bilayer indicated. The side chains of labeled residues are shown. The boxed area highlights the location of the Ca^2+^-clasp. **b**, Density for the Ca^2+^-clasp is missing in the absence of Ca^2+^. The structure of the Ca^2+^-clasp region that is observed in the Ca^2+^-bound closed structure (green) is shown in comparison with the cryo-EM density in this region in the Ca^2+^-free map, showing that the density for the Ca^2+^ ion and surrounding protein residues are missing in the absence of Ca^2+^. Ca^2+^ is depicted as a green sphere and two aspartate residues that coordinate Ca^2+^ as part of the Ca^2+^ clasp are shown as sticks. **c**, Refined cryo-EM maps of two conformations (blue, gray) of Ca^2+-^ free BEST1_345_ that were identified using 3D classification are depicted. The cryo-EM maps are aligned according to their membrane-spanning regions, with the relative rotation between the cytosolic regions indicated.

**Fig. S9.**
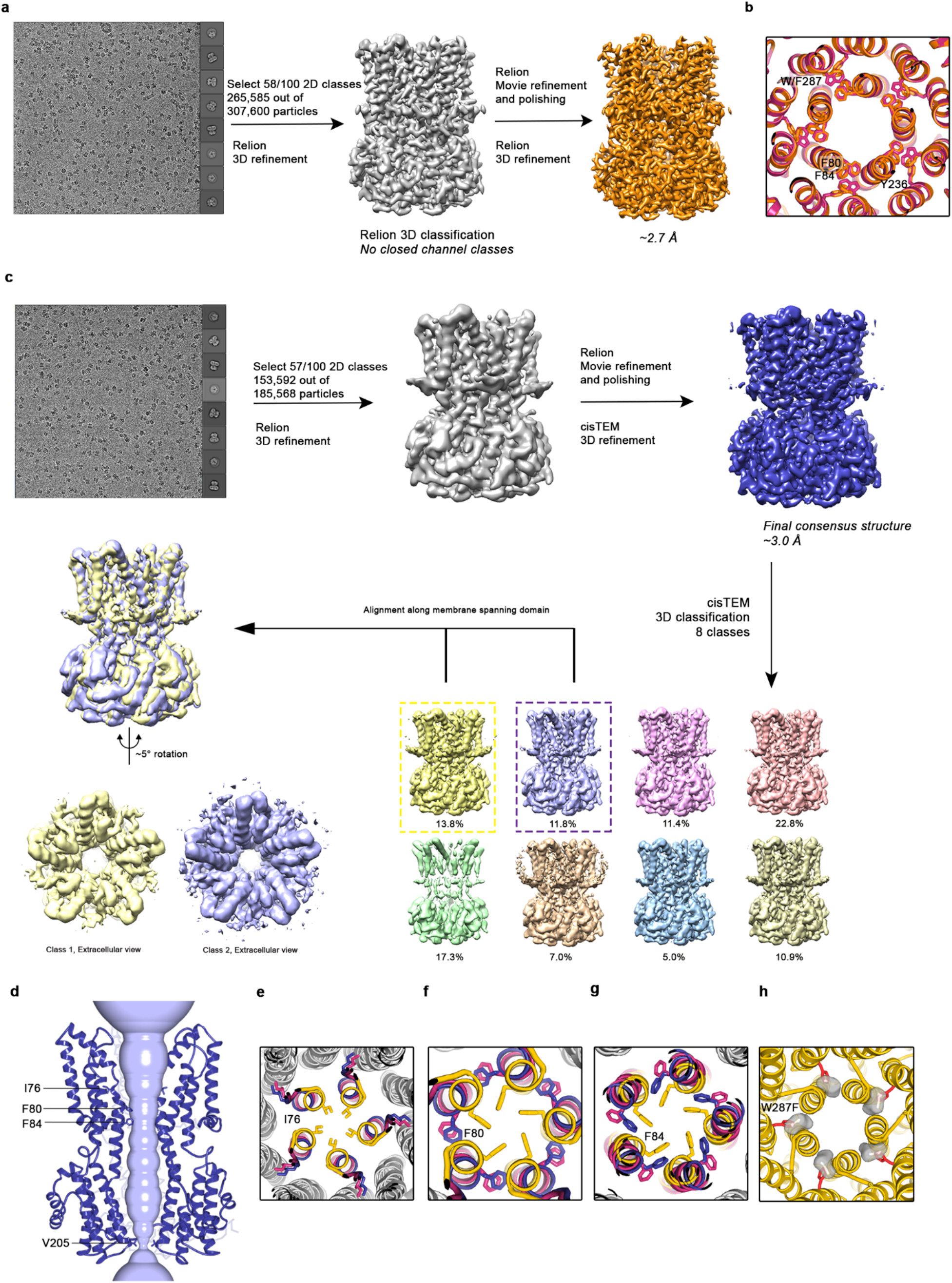
Cryo-EM workflow for Ca^2+^-bound and Ca^2+^-free BEST1_345_ W287F datasets. **a**, Cryo-EM workflow for Ca^2+^-bound BEST1_345_ W287F. A detailed description can be found in Methods. **b**, Comparison of F80, F84, W287 (or the W287F mutation), and Y236 between the open (pink; BEST1_345_) and the Ca^2+^-bound W287F mutant (orange; BEST1_345_ W287F) structures. A cutaway view is shown from the extracellular perspective. **c**, Cryo-EM workflow for Ca^2+^-free BEST1_345_ W287F. A detailed description can be found in Methods. **d**, Even in the absence of Ca^2+^, the neck of the W287F mutant is open. The minimal radial distance from the center of the pore to the nearest van der Waals protein contact is shown as a light blue surface. Two subunits of Ca^2+^-free BEST1_345_ W287F are depicted as ribbons; three are omitted for clarity. Amino acids in the neck an aperture regions are drawn as sticks. Approximate boundaries of the lipid membrane are indicated. **e-g**, Comparison of neck-lining residues I76 (e), F80 (f) and F84 (h) between Ca^2+^-free BEST1_345_ (yellow), Ca^2+^-free BEST1_345_ W287F (blue) and the open conformation of Ca^2+^-bound BEST1_345_ (pink). Side chains of labeled residues are depicted in each panel, viewed as a cutaway from the extracellular space. Helices are represented as ribbons and those not lining the pore are colored in gray. **h**, Modeling of a phenylalanine residue (red sticks) in place of W287 in the closed structure of BEST1_345_ (yellow ribbons) introduces a void (gray surface) behind the neck. The void was identified and displayed using Pymol (pymol.org).

**Fig. S10.**
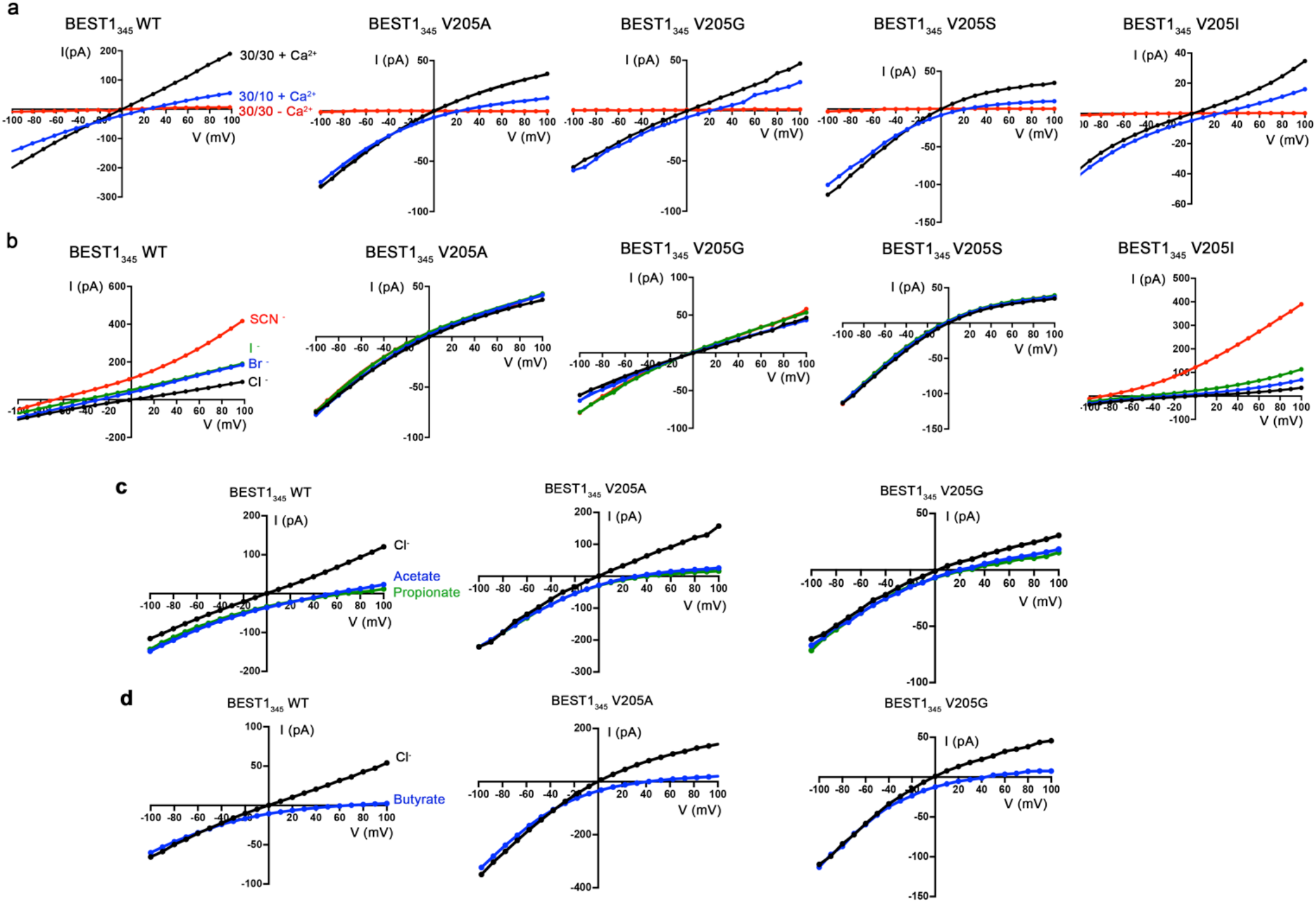
Current-voltage relationships of aperture mutants. **a**, All aperture mutants exhibit indistinguishable relative permeabilities of Cl^−^ versus K^+^ in comparison wild-type BEST1 and are Ca^2+^-dependent. Representative *I-V* relationships are shown for voltages stepped from −100 to +100 mV for the indicated standard conditions [cis/trans KCl concentration in mM, and ~ 300 nM [Ca^2+^]_free_ (+Ca^2+^) or 10 mM EGTA (-Ca^2+^)]. **b**, *I-V* relationships of BEST1_345_ WT (wild type) and BEST1_345_ V205 mutants that were used to determine permeabilities of Br^−^, I^−^, and SCN^−^ relative to Cl^−^. After first recording using symmetric 30 mM KCl (black *I-V* trace), the solution on the *trans* side was replaced (by perfusion) with solutions containing 30 mM KBr (blue), KI (green), or KSCN (red). **c**, Representative *I-V* relationships that were used to determine permeabilities of acetate and propionate relative to Cl^−^. Experiments were performed as in (b) except that the solution on the *trans* side was replaced with a solution containing 30 mM KCH3COO (blue) or KC2H5COO (green). **d**, Representative *I-V* relationships that were used to determine permeabilities of butyrate relative to Cl^−^ for the indicated constructs. Experiments were performed analogously to those described in (b) but used sodium salts: after recording in symmetric 30 mM NaCl (black I-V trace) the solution on the *trans* side was replaced with solution containing 30 mM NaC_3_H_7_COO (blue).

**Movie S1. Opening transitions.** This movie shows a morph between the closed and open conformations. Depictions are as described in Fig. 2a, b.

